# Cell-free expression and SMA copolymer encapsulation of a functional receptor tyrosine kinase disease variant, FGFR3-TACC3

**DOI:** 10.1101/2024.06.04.596442

**Authors:** Alexander J D Snow, Tharushi Wijesiriwardena, Benjamin J Lane, Brendan Farrell, Polly C Dowdle, Matilda Katan, Stephen P Muench, Alexander L Breeze

## Abstract

Despite their high clinical relevance, obtaining structural and biophysical data on transmembrane proteins has been hindered by challenges involved in their expression and extraction in a homogeneous, functionally-active form. The inherent enzymatic activity of receptor tyrosine kinases (RTKs) presents additional challenges. Oncogenic fusions of RTKs with heterologous partners represent a particularly difficult-to-express protein subtype due to their high flexibility, aggregation propensity and the lack of a known method for extraction within the native lipid environment. One such protein is the fibroblast growth factor receptor 3 fused with transforming acidic coiled-coil-containing protein 3 (FGFR3-TACC3), which has failed to express to sufficient quality or functionality in traditional expression systems. Cell-free protein expression (CFPE) is a burgeoning arm of synthetic biology, enabling the rapid and efficient generation of recombinant proteins. This platform is characterised by utilising an optimised solution of cellular machinery to facilitate protein synthesis *in vitro*. In doing so, CFPE can act as a surrogate system for a range of proteins that are otherwise difficult to express through traditional host cell-based approaches. Here, functional FGFR3-TACC3 was expressed through a novel cell-free expression system in under 48 hours. The resultant protein was reconstituted using SMA copolymers with a specific yield of 300 µg/mL of lysate. Functionally, the protein demonstrated significant kinase domain phosphorylation (*t<0*.*0001*). Currently, there is no published, high-resolution structure of any full-length RTK. These findings form a promising foundation for future research on oncogenic RTKs and the application of cell-free systems for synthesising functional membrane proteins.

## Introduction

Integral membrane proteins (IMPs) are simultaneously overrepresented as drug targets and underrepresented in the Protein Data Bank (PDB) compared to their soluble counterparts [1, 2]. The structural study of IMPs is often complicated by the difficulty inherent in producing and isolating them [3]. Many IMPs serve as potential drug targets that have not yet been structurally characterised, as most attempts using cell culture-based expression have produced inactive or degraded protein of insufficient quality for study.

Historically, cell-free protein expression (CFPE) systems have largely been based on *E. coli* lysate and used where the target protein would be toxic to the cellular systems traditionally used to express it [4, 5]. However, these methods were typically associated with expensive methodology, reducing their utility in large-scale reactions; poor specific yields and an inability to recreate native eukaryotic post-translational modifications (PTMs) present further difficulties [6]. Attempts to overcome these limitations have begun to show progress within the last decade. Eukaryotic lysates can include machinery capable of recapitulating native-like PTMs and yields have been increased thanks to the adoption of T7-polymerase systems to allow coupled transcription-translation within the same system [6]. Notably, a CFPE system based on *Nicotiana tabacum* BY-2 lysate is capable of PTM incorporation, high specific yields, and can be effectively scaled between the microlitre and litre ranges [7, 8]. Such systems allow the expression of eYFP at up to 3 mg/ml lysate used over a 48-hour synthesis [8]. The maturation of CFPE systems as a technology has culminated in the production of traditionally difficult IMPs such as epidermal growth factor receptors (EGFRs) in an active form [9].

Typically, for IMPs the requirement for a lipidic environment means they are isolated by encapsulating the hydrophobic transmembrane region in a detergent micelle, typically n-dodecyl-β-D-maltopyranoside (DDM) or similar [3]. This can often solubilise and stabilise the membrane protein but strips away the membrane lipids around it, precluding the study of the protein in native or native-like environments.

This membrane environment can be re-introduced through reconstitution into lipid nanodiscs, micelles or bicelles. However, this generally requires the addition of formulated lipid mixtures, that are unlikely to match the exact environment of the protein *in vivo [10]*. The introduction of synthetic copolymers such as styrene and diisobutylene-maleic acid (SMA, DIBMA) to the field presents a step forward, wherein an IMP can be directly removed from the membrane while retaining local lipids [11, 12]. Where post-synthesis solubilisation has proven ineffective, some CFPEs allow membrane protein insertion into lipid microsomes or even protein-lipid nanodiscs, which eliminates the need for co-expressed nanolipoproteins as used with the receptor tyrosine kinase (RTK) EGFR [8, 13, 14].

As a family of integral-membrane protein kinases, RTKs are responsible for propagating a wide variety of signals across the plasma membrane [15, 16]. RTKs can be further classified into 20 subfamilies, which share a general arrangement consisting of an N-terminal extracellular domain (ECD), a single pass transmembrane *α*-helix, and a C-terminal, intracellular tyrosine kinase domain [15]. While this arrangement is conserved across all subfamilies, the composition of the ECD, the number of kinase domains present, and the activation mechanisms vary significantly across them. For example, Axl and Eph receptors use a bidentate ligand to directly induce dimer formation while EGFR undergoes ligand-induced conformation changes to undergo lateral dimerisation and subsequent activation [17, 18].

One family of RTKs is the fibroblast growth factor receptors (FGFRs), which among its four variants (FGFR1-4) and their associated splice-variants can become activated via a variety of mechanisms depending on the combination of receptor and ligand (fibroblast growth factor, FGF). One of 18 mammalian FGFs can bind the receptor ECD between two of the three Ig-fold domains (D2, 3) that comprise it, in a 1:1 stoichiometry [19]. Paracrine signalling also requires the recruitment of heparan sulphate from the local extracellular matrix while endocrine signalling is characterised by *α*/β-Klotho recruitment [20-22]. This ECD:FGF:cofactor complex disrupts the auto-inhibitory role of the D1 domain allowing dimer formation and receptor activation. It is currently theorised that the specific ligand/cofactor combination influences the juxta-membrane spacing of the ECD, and by extension the angle of the transmembrane *α*-helix pair [23, 24]. In all cases, downstream effects are modulated by the *trans*-autophosphorylation of the C-terminal tyrosine kinase domains, which form an asymmetric, active dimer in which the activation loop of one domain protrudes into the nucleotide-binding region of the counterpart [25]. Tyrosine phosphorylation then proceeds within the loop before becoming more widespread [26]. Current theories for the disparate effects caused by FGFR activation heavily feature the angle at which the transmembrane helices of an active dimer cross, a feature likely induced by changes in D2-D3 domain angle with different ligands [24].

Dysregulation of FGFRs is a major cause of disease, with all four FGFRs featuring oncogenic point mutations across all four domains [27]. These generally affect signalling through either the promotion of a ligand-independent dimer or a constitutively active kinase domain [16, 28]. An FGFR mutation of note is a fusion with transforming acidic coiled-coil containing protein 3 (TACC3), producing FGFR3-TACC3 through a tandem duplication event [29, 30]. FGFR3-TACC3 features a C-terminal coiled-coil domain as well as constitutive phosphorylation of activation loop tyrosine residues, thus causing aberrant signalling [30, 31]. FGFR3-TACC3 is often implicated in the initiation and progression of glioblastoma, oral, gallbladder, urinary bladder, and cervical cancers, among others [29, 32].

Given the signalling dysregulation caused by FGFR3-TACC3, obtaining high-resolution structural insights would hold high value in the search for superior therapeutic interventions. Structural study of full-length RTKs has so far been limited to low-resolution or partially-resolved views, and no such data has been published for any FGFR [33]. Biophysical characterization of this important oncoprotein has hitherto been confounded by difficulties in producing and isolating it while retaining activity. This effectively limits the study of full-length protein to experiments that do not require purified sample [22], thus precluding high-resolution structural analysis. Hence, it was considered an ideal target for a CFPE and copolymer solubilisation approach. Here, we report the successful cell-free synthesis, purification and encapsulation of FGFR3-TACC3 in SMA lipid particle (SMALP) nanodiscs. We confirm the product is constitutively active as expected, as well as capable of binding the canonical ligand FGF1. These data present a substantial development in our ability to structurally characterise a previously intractable class of disease-causing proteins.

## Results

### Construct design

*In vitro* and *in vivo* studies in bladder cancer have demonstrated that the FGFR3-TACC3 variant designated RT112 (after the bladder cancer cell line in which it was identified) is not only clinically relevant but that it also expresses a range of cellular behaviours that distinguish it from canonical FGFRs and imply distinct activation mechanisms [31]. Difficulties in expressing and solubilising the protein in sufficient quantities for downstream biophysical analyses have thus far hindered progress in unravelling the basis for its aberrant activity. Therefore, we attempted protein production using the ALiCE (LenioBio GmbH) CFPE system, which is natively capable of inserting transmembrane proteins into membrane compartments thought to resemble those of *N. tabacum* early endosomes [8].

Cell-free expression typically requires the incorporation of the target gene into a specialised vector, with the system itself engineered to take advantage of a T7 polymerase [34]. To support microsomal membrane insertion, RT112 was cloned into the pALiCE02 vector. The construct also included a C-terminal, HRV-3C-cleavable GFP reporter protein and 10x His tag to allow efficient detection of synthesized protein and subsequent bulk purification.

## Protein production

After the addition of the RT112-coding construct in pALiCE02 to 50µl ALiCE lysate, fractions were monitored for an increase in GFP fluorescence which would correlate with the presence of microsome-bound RTT112. GFP fluorescence could be detected after 2 hours, and the intensity increased in an approximately linear fashion to 60% of the maximum over the first 10 hours, after which it started to plateau (Figure 1a). The observed fluorescence only increases by 20% in the final 24 hours of the reaction (Figure 1a). While degradation of the protein is observed, it is confined to the non-membraneous fraction (supplementary figure 8). Reconstitution of RT112 can be achieved in DDM micelles, and the produced protein can be partially purified through IMAC and size-exclusion chromatography. However, solution Mw analysis by SEC, SDS-PAGE (Supplementary Figure 1) and native-PAGE both show a wide variation, as well as some that may represent dimeric protein (unpublished observations). This, combined with the strong likelihood that physiologically relevant protein-lipid interactions made by the RT112 transmembrane domain are not retained in DDM micelles, likely renders the method a sub-optimal choice for functional studies. Therefore, subsequent attempts aimed to capture RT112 with the surrounding lipid environment in protein-SMALPs. Addition of 2.5% w/v SMA2000 resulted in a notable increase in sample clarity over 2.5 hours. The soluble fraction was purified by IMAC and size-exclusion chromatography,.

**Figure 1:**
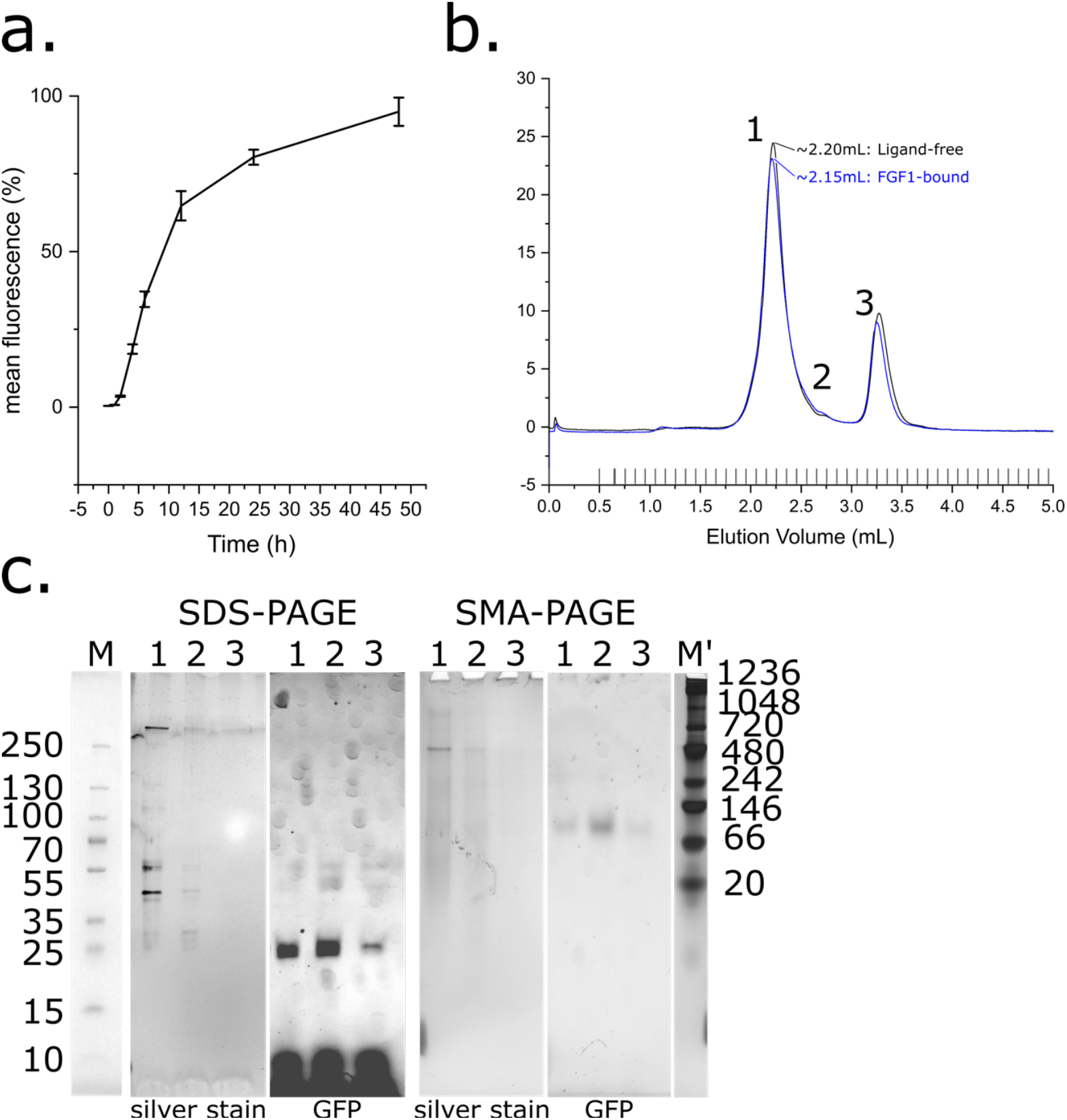
Cell-free expression and purification of RT112 in SMALPs. (a) Fluorescence normalised against the maximum observed with time in hours. Error bars represent the standard error across 3 repeats. (b) Size-exclusion chromatogram of RT112. Black trace represents ligand-free protein, blue represents ligand-bound (in the presence of FGF1). Peaks 1, 2 and 3 eluted at approx. 2.20, 2.70 and 3.30 mL respectively. (c) PAGE analysis of fractions collected during size-exclusion chromatography of RT112. Fractions 1,2 and 3 from b. were analysed by SDS and SMA-PAGE, and visualised using silver stain and in-gel GFP fluorescence. Denatured ladder (M) was obtained from the SDS-PAGE pre-silver-stain and aligned to SDS-PAGEs. Native ladder (M’) was obtained post silver stain and aligned to SMA-PAGEs.

Incubation of a pre-SEC sample with FGF1 produced a small increase in molecular weight, though due to the small size of the ligand (18 kDa) this difference was not well resolved (Figure 1b).SDS-PAGE of the ligand-free SEC fractions resulted in fluorescent bands, strongly implying the purified species is RT112. The Mw of these species was lower than expected on SDS-PAGE, falling into a range of masses between 100 kDa to 25 kDa. The same fractions on SMA-PAGE [35] inhabit a single band at approximately 130 kDa, corresponding well to the sequence-derived Mw of 136 kDa (Figure 1c, Supplementary Table 1). This suggests the SDS-PAGE result does not reflect protein degradation but may perhaps, arise due to residual lipid and polymer still adducted after denaturation in SDS sample buffer Overall, RT112 purified through these methods has a specific yield of approximately 300 µg per 1mL lysate used (Supplementary Figure 5).

### Kinase Activity

RT112 features constitutive kinase activity *in vivo*, so to test whether the protein produced through ALiCE was physiologically relevant, its kinase domain activity was assayed. An ADP-Glo functional assay was selected due to its sensitivity in detecting relatively few phosphorylation events resulting from trans-autophosphorylation, when compared to kinases with activity against heterologous substrates. Assays performed on RT112 previously treated with lambda protein phosphatase (LPP) to hydrolyse adventitious tyrosine phosphorylation showed a statistically significant (P<0.0001) increase in luminescence compared to untreated RT112. As such, ATP turnover in the sample was indicative of activity that was present at any point post-treatment (Figure 2). These data are consistent with the RT112 produced in this study being catalytically active in a ligand-independent manner, in agreement with behaviour *in vivo*. To rule out the possibility of observed activity arising from another, co-purified kinase, the same assay was performed on un-transfected ALiCE lysate, purified with precisely the same method as RT112. This un-transfected sample showed no activity (Figure 2). We were unable to further compare the activity of our ALiCE-generated RT112 with that derived via more conventional methods because, in our hands, the protein fails to express and purify to a sufficient yield for study in all tested cell-based expression systems (data not shown).

**Figure 2:**
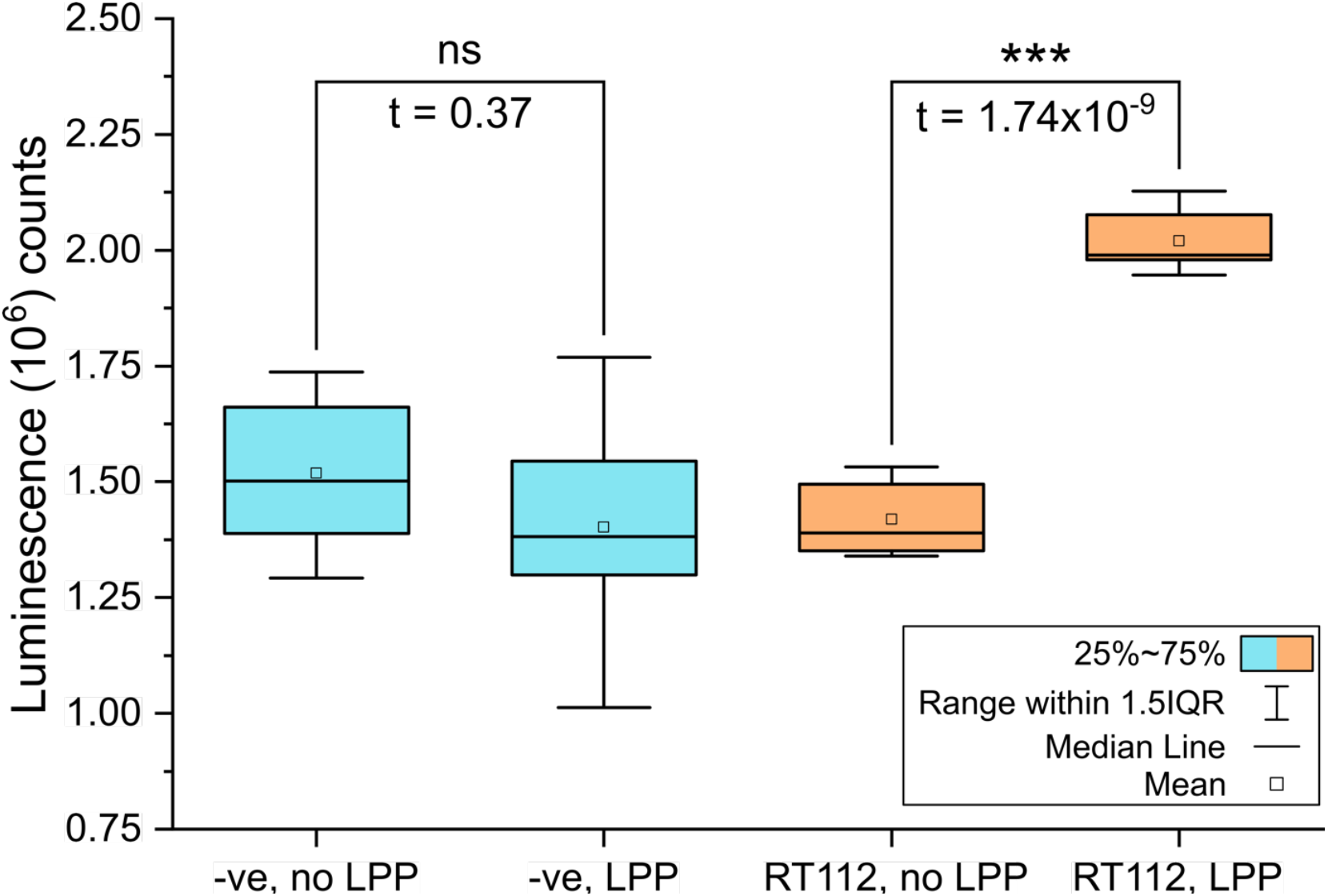
Kinase activity of RT112. Box plot of ADP-glo results of un-transfected, purified lysate (red) and RT112-SMALPs (orange) in an untreated and lambda-phosphatase (LPP)-treated state. Statistical significance was determined using a twinned t-test between the mean counts for each dataset.

### Growth Factor Binding

To establish whether our ALiCE-expressed RT112 protein retained ligand-binding capability, we incubated it with FGF1. The small increase in solution Mw (36 kDa) this would represent was not readily apparent on SEC; however a clear band deflection was observed on SMA-PAGE (Figure 1b, Figure 2a). FGF1 incubation also resulted in an additional species weighing approximately 450 kDa, visible both through SEC and SMA-PAGE (Figure 3a,b) [35]. The presence of an RT112.FGF1 complex was confirmed through mass photometry. FGF1 incubation induces a clear shift in particle size wherein the predominant population increases from 126 kDa to 242 kDa, strongly suggestive of a concomitant dimerisation event (Figure 3c). A small population of monomeric protein remained despite the 10x molar excess of FGF1 used, perhaps suggesting the ligand does not fully stabilise an RT112 dimer. The larger 480 kDa species observed on SMA-PAGE was not visible through mass photometry, perhaps implying that it is not a relevant solution state.

**Figure 3:**
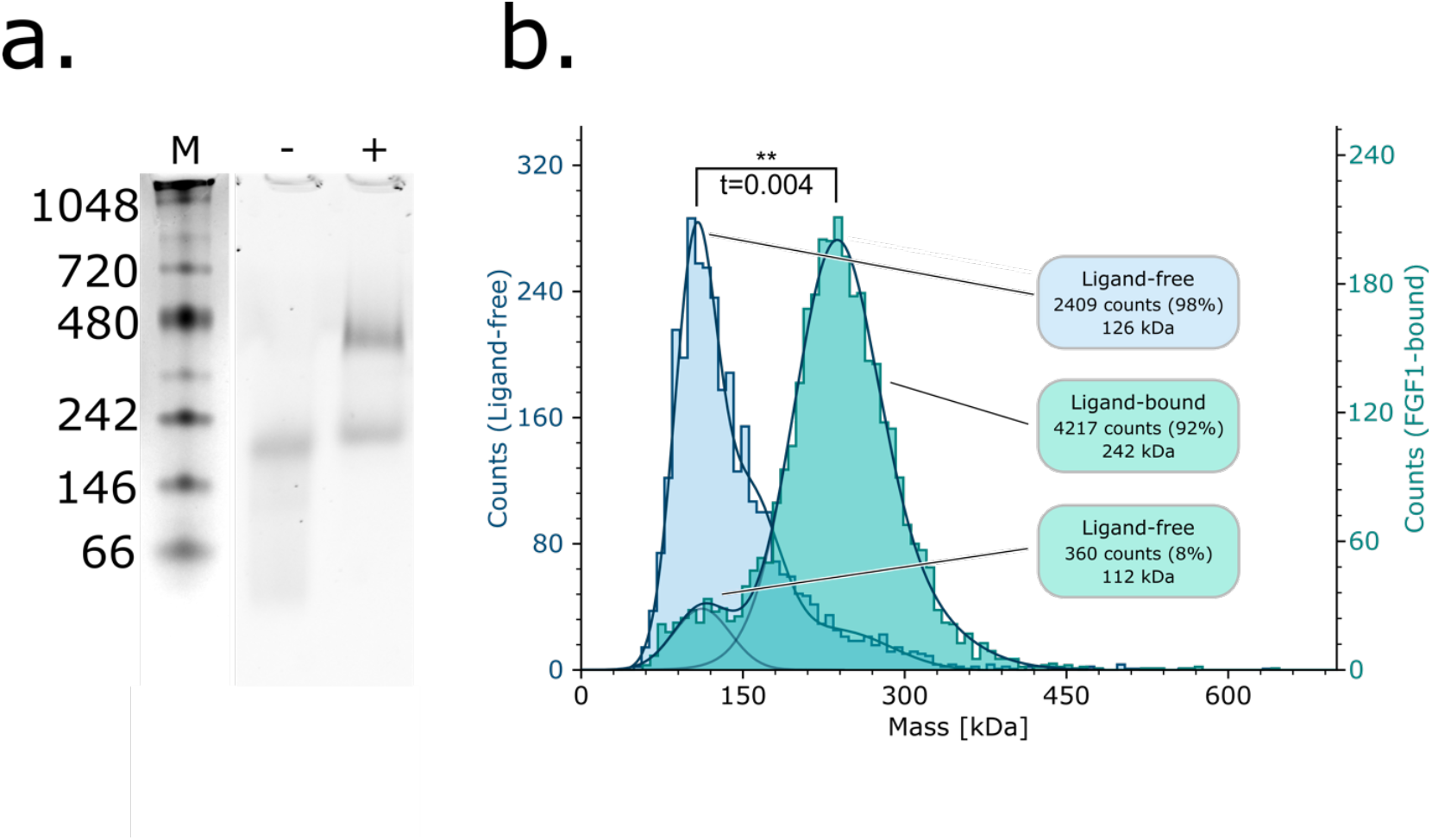
RT112 shows growth factor binding activity. (a) Native-PAGE of RT112 in a ligand-free (-) and FGF1-incubated (+) state. Samples are shown under GFP-fluorescence. The ladder (M) was composited from a visible image of the same gel. (b) Mass photometry histograms of ligand-free (blue) and FGF1-incubated (cyan) RT112. Major peaks from both histograms are labelled in the respective colours, and counts are on the left and right respectively. Statistical significance was determined using a twinned t-test between the mean counts for each dataset.

## Discussion

To date, the number of publications featuring preparative-scale production of intact, functional RTKs remains low, in part due to the difficulty in producing these proteins through traditional methods. Cell-free systems have been used previously to achieve functional RTK expression, but this required *E. coli* codon optimisation across the entire sequence and co-expression with nanolipoproteins to allow for membrane insertion [9]. Our work with RT112 builds on earlier studies to show that RTK disease variants can be produced using native codon bias and no further modification of the synthesis system. The resulting protein can be reconstituted into SMALPs, retaining the native-like lipid environment, whilst attaining a high degree of purity (>90%). RT112 clearly expresses ligand-independent kinase activity, as expected from a constitutively active RTK oncoprotein. Our data also confirm that the dimerisation propensity of RT112 is more complex than other TACC3 fusions, which often form constitutive dimers [32]. Instead, RT112 dimerisation becomes more favourable with FGF1 binding, though this remains transient as, through both SMA-PAGE and mass photometry, populations corresponding to a monomer can still be observed. SMA-PAGE shows these monomers still have a ligand bound.

In summary, we demonstrate here that an oncogenic FGFR3-TACC3 fusion can be effectively produced and reconstituted into SMALPs, then purified at preparative scale, using a cell-free protein expression system. RT112 thus prepared not only retains constitutively active kinase domain activity, but also undergoes dimerisation in the same manner as natively-sourced protein. From these data we propose that CFPE is a highly advantageous method for exploring these challenging targets and can provide a flexible platform for further biophysical and high-resolution structural characterisation.

## Acknowledgements

This work was supported by the Medical Research Council (MR/W000369/1) and the Biotechnology and Biological Sciences Research Council (BB/W008017/1). We would like to thank Dr. Tom Bunney (University College London) for his advice and help throughout this work as well as Dr. Yvonne Nyathi and Urjaa Kulkarni (University of Leeds) for their contributions. Dr. Arnout Kalverda (Astbury Biostructure Laboratory NMR Facility) is thanked for expert technical support for the NMR experiments.

## Author Contributions

A.L.B and S.P.M conceived the project. B.F designed the gene construct and enacted preparative mutagenesis. A.J.D.S and T.W expressed, solubilised and purified RT112, and carried out all characterisation experiments and processing of these. A.J.D.S designed the plasmid for FGF1. A.J.D.S and P.C.D over-expressed and purified FGF1. P.C.D carried out ^1^H-NMR of FGF1. B.J.L assisted with all laboratory-based stages. All authors contributed to the writing of the manuscript.

## Data Availability

All data generated or analysed during this study are included in this published article (and its Supplementary Information files).

## Materials and Methods

### Construct design

The amino acid sequences for all proteins produced in this study are recorded in Supplementary Table 1. The FGFR3-TACC3 RT112 used in this study was based upon a variant of wildtype FGFR3 IIIb. To improve stability during synthesis, all tyrosine phosphorylation sites on the kinase domain were mutated to phenylalanine except Y647, which is essential for activity. This Y577F/Y648F/Y724F/Y760F/Y770F mutation series is denoted ‘5F1Y’. Additional C228R, S249C and Y375C mutations were introduced to the ECD along with the constitutively activating K652E kinase-domain mutation, seen in clinically occurring FGFR3-TACC3 RT112. A final construct was produced from synthetic GeneArt^®^ Strings^™^ (Thermo Fisher Scientific) including a C-terminal, HRV-3C cleavable sfGFP; a 10x His tag wasincluded for ease of purification. Finally, this was cloned into pALiCE02 using In-fusion cloning (Takara). Amplification of FGFR3-TACC3.RT112.pALiCE02 involved transformation into One Shot^™^ OmniMAX^™^ 2 T1 R competent cells (Thermo) by the manufacturer’s method, then liquid-culture growth to a 1 L scale in LB+100 µg/mL ampicillin. Finally, DNA was extracted and purified using a Qiagen Plasmid maxiprep kit, eluting DNA into ultrapure water (Thermo) but otherwise using manufacturer instructions. Final DNA purity was assessed by running on a 2% agarose/TBE gel + 0.01% SYBRsafe, run at 110 V.

The gene for FGF1 was codon-optimised for *E. coli* expression and ordered in pET28a (Genscript).

### Cell-free protein synthesis

A 1 mL aliquot of ALiCE (LenioBio GmbH) cell-free reagent was thawed from -80°C in a room-temperature water bath, then centrifuged at 500g, 4 seconds to collect material at the bottom of the tube. Depending on experiment, either 50 or 60 µL of this was added to each of 16 perforated-top 2 mL tubes. To these FGFR3-TACC3.RTT112.pALiCE02 was added to a final 24 ng/µL concentration. The reaction then progressed for 48 hours, 23°C with 700 rpm orbital shaking using a ThermoMixer C (Eppendorf). Protein expression was assayed fluorometrically using a HIDEX plate reader, using a 485 nm excitation and 535 nm emission wavelength with one flash on low lamp power. These measurements were made in a 384-well white-bottom plate using 3x 20 µL samples per timepoint.

### Solubilisation and purification of FGFR3-TACC3

Post reaction, ALiCE lysate was pooled and centrifuged at 16,000 xg, 40 minutes, 4 °C to pellet organelles and microsomes. These were resuspended in 50 mM HEPES, 500 mM NaCl, 5% v/v glycerol, 1 mM TCEP, 1 mM MgCl_2_ 1 mM ATP, 1 mM EDTA pH 7.9. Additionally, 1 mM PMSF, 10 U DNAse I, 1 mini tablet EDTA-free protease inhibitor (Roche) were added per 10 mL buffer. For SMA solubilisation, 2.5% w/v SMA2000 was added and solubilisation progressed at 19 °C on a roller, 60 rpm for 2.5 hours. This SMA was hydrolysed in-house prior to use [36]. For DDM solubilisation 0.75% w/v DDM was added and the solubilisation progressed on a roller, 60 rpm at 4 °C for 45 minutes, after which a further 0.75% w/v DDM was added and the solubilisation progressed another 45 minutes. In both cases, insoluble material was pelleted at 130,000 xg, 4 °C for 1 hour. The soluble fraction was then loaded to 0.5 mL Ni-FF resin (Cytiva) pre-equilibrated into 50mM HEPES, 500 mM NaCl, 1 mM TCEP, 1 mM MgCl_2_, 10 mM imidazole pH 7.9 via 5 CV ultrapure water (Thermo). This was incubated at 4 °C on a roller, 60 rpm for 1 hour then loaded to a gravity column. The resin was washed with 10 CV of the buffer above, then His-tagged protein was eluted using 50 mM HEPES, 500 mM NaCl, 1mM TCEP, 1 mM MgCl_2_, 500 mM imidazole pH 7.9. Protein-containing fractions were pooled and concentrated using a 50 kDa MWCO spin concentrator (Sartorious) at 4500 xg, 4 °C until the total volume equalled 200 µL or less. This was then loaded to a Superose 6 Increase 5/150 GL column pre-equilibrated in 50 mM HEPES 500 mM NaCl 0.5% v/v glycerol 1 mM TCEP, pH 7.9 using a sample loop. Protein-containing fractions were used immediately and incubated on ice unless an assay required otherwise.

Purification fractions were visualised on a 4-20% TGX SDS-PAGE (Bio-Rad), run at 200 V, 35 minutes with TGX SDS-PAGE buffer (Thermo). In all cases sample was denatured through addition of SDS-PAGE loading dye concentrate (Cold Spring Harbor recipe) and incubation at 36 °C for 10 minutes immediately prior to loading. GFP fluorescence was detected using a 15-30s exposure on an iBright (Thermo) imager on an unstained gel. Gels were visualised either by Quick Coomassie (Thermo), or by silver staining, using a commercially available kit (Thermo) and following manufacturer instructions. All stained gels were imaged using a Bio-Rad GelDoc Go (Thermo). Native/SMA PAGEs were run using identical gels but with 1x TGX native PAGE buffer (Thermo). Here, a 10x molar excess of FGF1 was incubated with RT112 for 30 minutes at 4°C, and the result concentrated using a 50kDa MWCO concentrator (Sartorious) before SEC using the same method as ligand-free RT112. Samples were produced using a 1:1 ratio of native PAGE loading dye (CSH recipe) and sample, with no incubation time. This was run at 120 V and visualised identically to SDS-PAGE gels. All BCA protein concentration assays were performed using a commercial kit (Thermo) and following manufacturer-recommended protocols, in a 96-well plate format. The results were collected using a HIDEX plate reader measuring OD_586_.

### Expression and purification of FGF1

The gene for FGF1 was transformed into BL21(gold) competent cells (Sigma-Aldrich) using the supplier protocol and plated on LB-agar with 25 ug/mL kanamycin and grown at 37 °C over 18 hours. 25mL LB+25ug/mL kanamycin liquid cultures were produced from a single colony and grown at 37 °C, 200 rpm for 18 hours. Two 500 mL LB+25 ug/mL kanamycin liquid cultures were grown using 5mL of these as inoculant in baffled expression flasks at 37 °C 220rpm until OD_580_ = 0.7, then induced with 0.8 mM IPTG and grown for a further 18 hours at 18°C, 180 rpm. Cells were pelleted at 4000g, 4 °C for 20 minutes then snap-frozen in liquid nitrogen and stored at - 80C. When required, pellets were thawed and resuspended in 50 mM HEPES, 500 mM NaCl, 1mM TCEP, 0.5% w/v glycerol pH 7.8, with 1 mini EDTA-free protease inhibitor tablet (Roche) and 1mM PMSF added per 10mL buffer used. Cells were lysed using a disruptor at 25 kPSI, 5°C then lysate was clarified at 18,000 xg, 4 °C for 40 minutes. The soluble fraction was loaded to a 5 mL Ni-FF prepacked column over 1 hour then eluted using an imidazole gradient between 10 and 500 mM over 10 CV. Protein-containing fractions were pooled and concentrated at 4500 xg, 4°C using a 3 kDa MWCO spin concentrator (Sartorius) until the final volume was 0.5 mL. This was then loaded on to a Superdex 200 Increase 10/300 GL column pre-equilibrated in 50 mM HEPES, 500 mM NaCl, 0.5% v/v glycerol, 1 mM TCEP, pH 7.9 using a sample loop. The protein was estimated at >95% purity, with a total yield of >100 mg/L LB (Supplementary Figure S2). Finally pure FGF1 was concentrated to 150 µM using a 3 kDa MWCO 500 µL spin concentrator (Sartorius) then snap-frozen in liquid nitrogen and stored at -80°C in 200 µL fractions. The quality of FGF1 produced was assessed through 1D ^1^H-NMR (Supplementary Figure 2).

## 1D ^1^H-NMR

FGF1 samples were prepared by buffer-exchanging the original 150 µM stock into 50 mM sodium phosphate, 100 mM NaCl, 5% v/v D_2_O, pH 7.5 using a PD-10 desalting column, operated using the manufacturer’s instructions. The concentration of the eluted protein was then estimated using the A_280_ and a sample was diluted to 50 μM, 560 μL. This was then transferred to a 5mm NMR tube. The ^1^H spectra were recorded at 25°C using a Bruker Avance III HD spectrometer at 750 MHz with a 5 mm TCI cryoprobe and processed with TopSpin 3.2 (Bruker Biospin).

### ADP-Glo kinase assay

RT112 used in this assay was prepared identically but purified post-IMAC using a 1mL Q-HP column (Cytiva). To achieve this, post-IMAC fractions were buffer-exchanged into 50 mM HEPES, 150 mM NaCl, 1 mM TCEP, 0.5% v/v glycerol pH 7.9, then concentrated with 50 kDa MWCO 500 µL spin concentrators (Sartorious) at 7000 xg, 4°C until a volume of 200 µL was achieved. This was loaded to the column using an ÄKTA pure system (Cytiva) with a sample loop, and species separated using a gradient to 50 mM HEPES, 2 M NaCl, 1 mM TCEP, 0.5% v/v glycerol pH 7.9. Purified protein was buffer-exchanged into 50 mM HEPES, 100 mM NaCl, 2 mM DTT, 2 mM MnCl_2_, 20 mM MgCl_2_, 0.1 mg/mL BSA, pH 7.9 using a PD-10 desalting column, then concentrated to 20 µM. All reactions were carried out in a 384-well white-bottom plate. 100 U lambda phosphatase was then added to all test cases (buffer was added in controls) and the sample was incubated for 30 minutes, 30°C. Phosphatase activity was quenched by adding 1mM sodium orthovanadate, and kinase activity resumed by adding 2 mM ultrapure ATP and incubating at 15°C, 1 hour. ADP-glo reagent was then added and the mixture was incubated for 2 hours, 15°C. Finally, kinase detection reagent was added, and the mixture was incubated for 1 hour, RT. Luminosity was measured using a HIDEX plate reader with a 1 second IR cutoff and a 20s orbital shake, 300 rpm prior to detection. All statistical analysis was performed using Origin 2020. The un-transfected control was produced and processed identically to RT112, however with ultrapure water added in place of input plasmid stock

### Mass Photometry

All mass photometry data were acquired on a REFEYN OneMP instrument with isolation unit (Refeyn Ltd.), using a standard drop-dilution method. Microscope coverslips were cleaned prior to use by washing in 100% isopropanol, followed by 6 minutes ultrasonic cleaning in ultrapure water (Thermo). Slides were then dried using a room temperature air stream. All FGFR3 samples were prepared to a 500 nM concentration. FGFR3-FGF1 complexes were prepared by incubation on ice, 30minutes with FGF1 at a 10x molar ratio. In all cases, focus was attained using an 18 µl drop of 50 mM HEPES, 500 mM NaCl, 0.5% v/v glycerol, 1 mM TCEP, pH 7.9, to which 2 µl of the relevant sample was added to begin the measurement. Each measurement consisted of a 60s recording at room temperature using a 525 nm laser at 1000 fps, 21 nm/px using AcquireMP software. All analyses were conducted using DiscoverMP (Refeyn Ltd.). All statistical analysis was performed using Origin 2020. Results were obtained in triplicate.

