## supplementary information for "Cell-free expression and SMA copolymer encapsulation of a functional receptor tyrosine kinase disease variant, FGFR3-TACC3"

**Keywords**
Cell-free, FGFR, RTK, oncoprotein, kinase, SMA

**Authors**
Snow, Alexander J D[1]; Wijesiriwardena, Tharushi[1]; Lane, Benjamin J[1]; Farrell, Brendan[1]; Dowdle, Polly C[1]; Katan, Matilda[2]; Muench, Stephen P[1,*] and Breeze, Alexander L[1,*]

[1]Astbury Centre for Structural Molecular Biology, University of Leeds, Leeds, LS2 9JT, UK
[2]Institute of Structural and Molecular Biology, Division of Biosciences, University College London, London, WC1E 6BT, UK

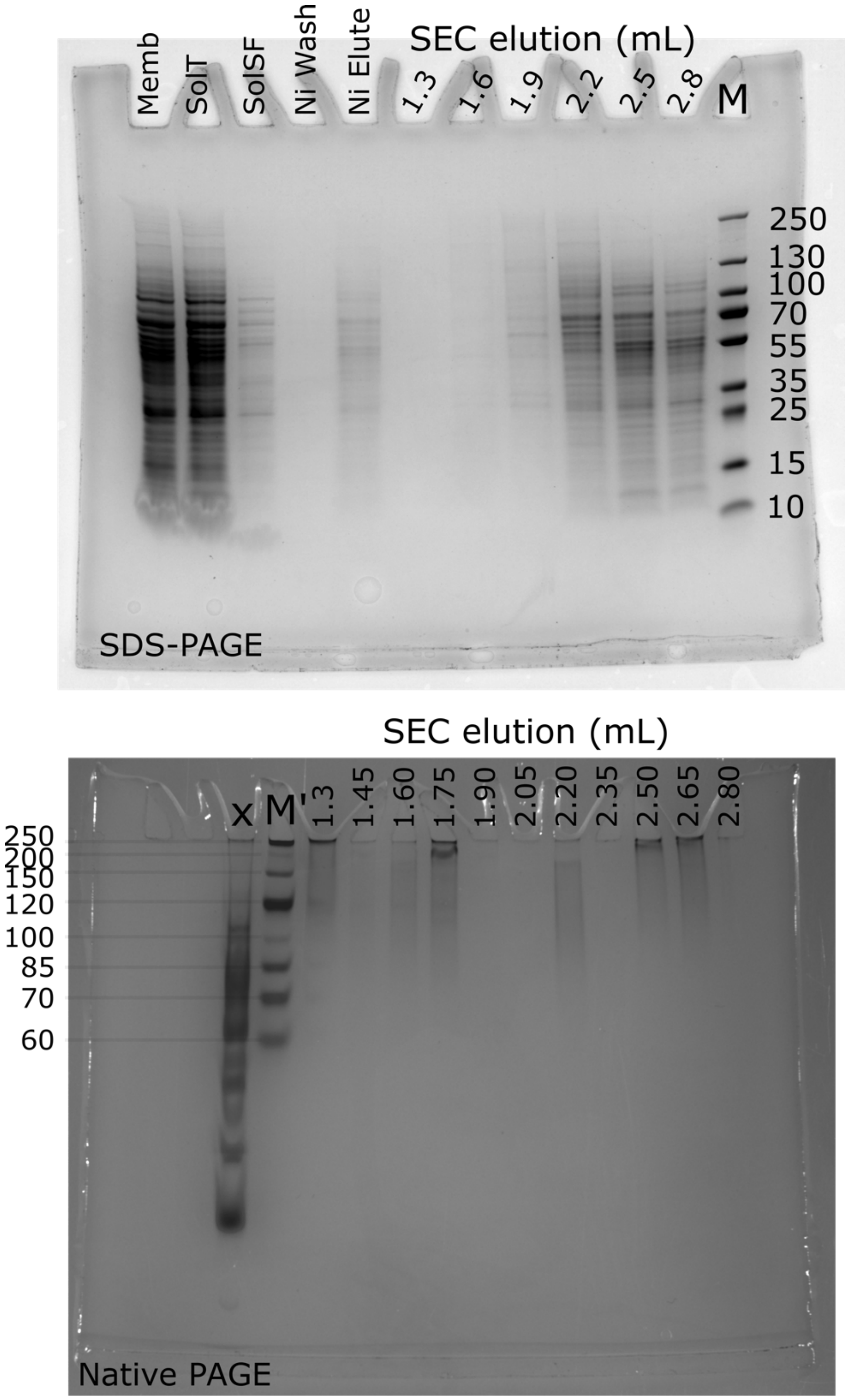


Figure S1: RT112 purification using DDM. Top, SDS-PAGE. Left to right: homogenized membrane fraction, DDM-treated membrane fraction, soluble fraction post-ultracentrifugation, nickel affinity wash and elution, and elution fractions from a superose 6 increase 5/150gl SEC column at volumes between 1.3 and 2.8 mL, pageruler prestained ladder. Bottom, native-PAGE. Left to right: high-Mw protein ladder (indicative), SEC elutions from 1.3 to 2.8 mL. X represents a lane unrelated to this work. Gels were Coomassie-stained.

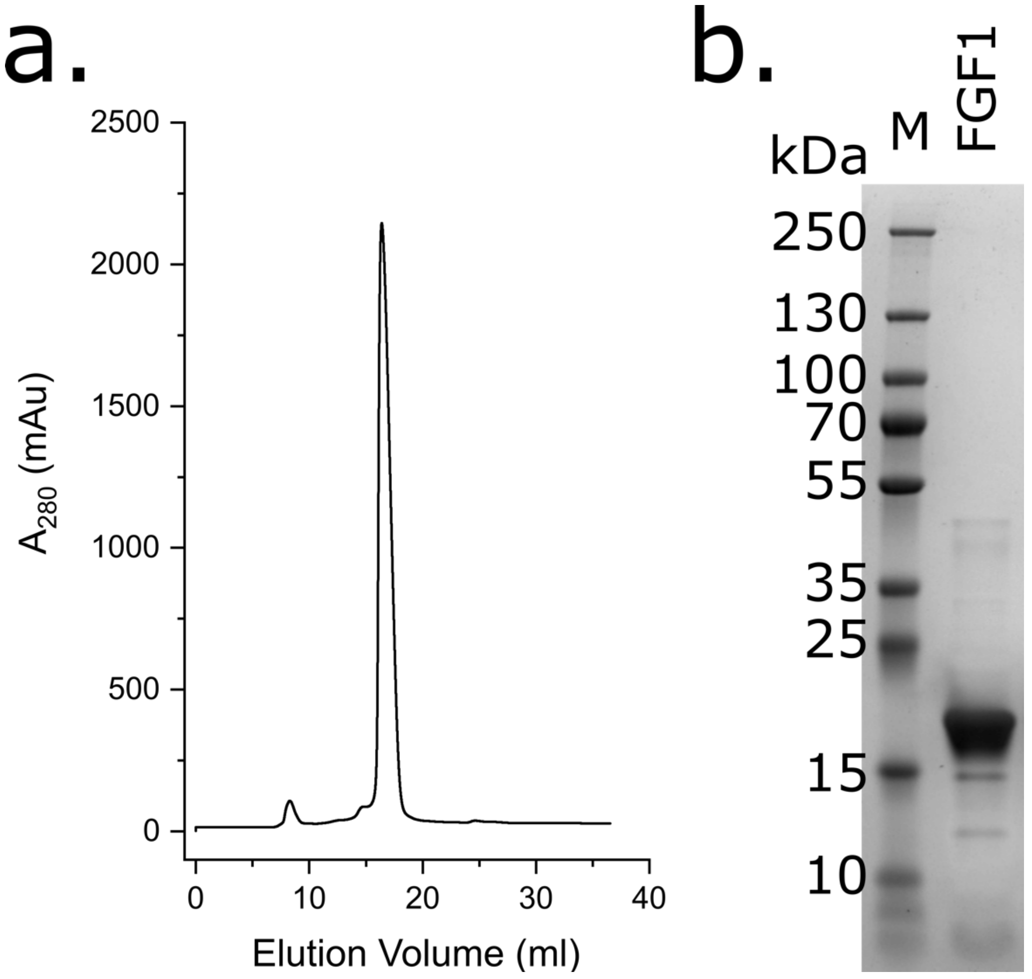


Figure S2: *Purification of FGF1:* (a.) Size-exclusion chromatography trace of FGF1. The peak at approx. 17ml corresponds to monomeric FGF1. (b.) SDS-PAGE of purified FGF1. The overloaded bands at 10-20 kDa correspond to the protein at >95% purity. An estimated 100mg per 1L LB were produced overall. SDS-PAGE has been cropped for clarity.

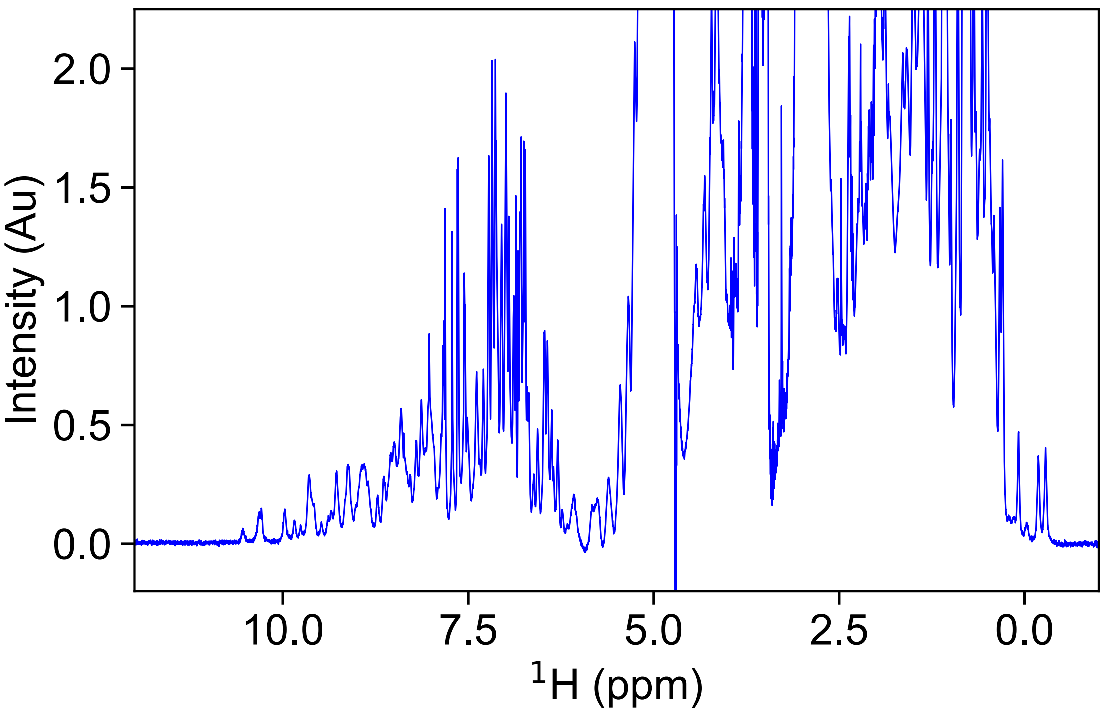


Figure S3: *1D ¹H-NMR spectrum of FGF1*. The extensive dispersion and good peak resolution is consistent with a correctly folded, monomeric protein.

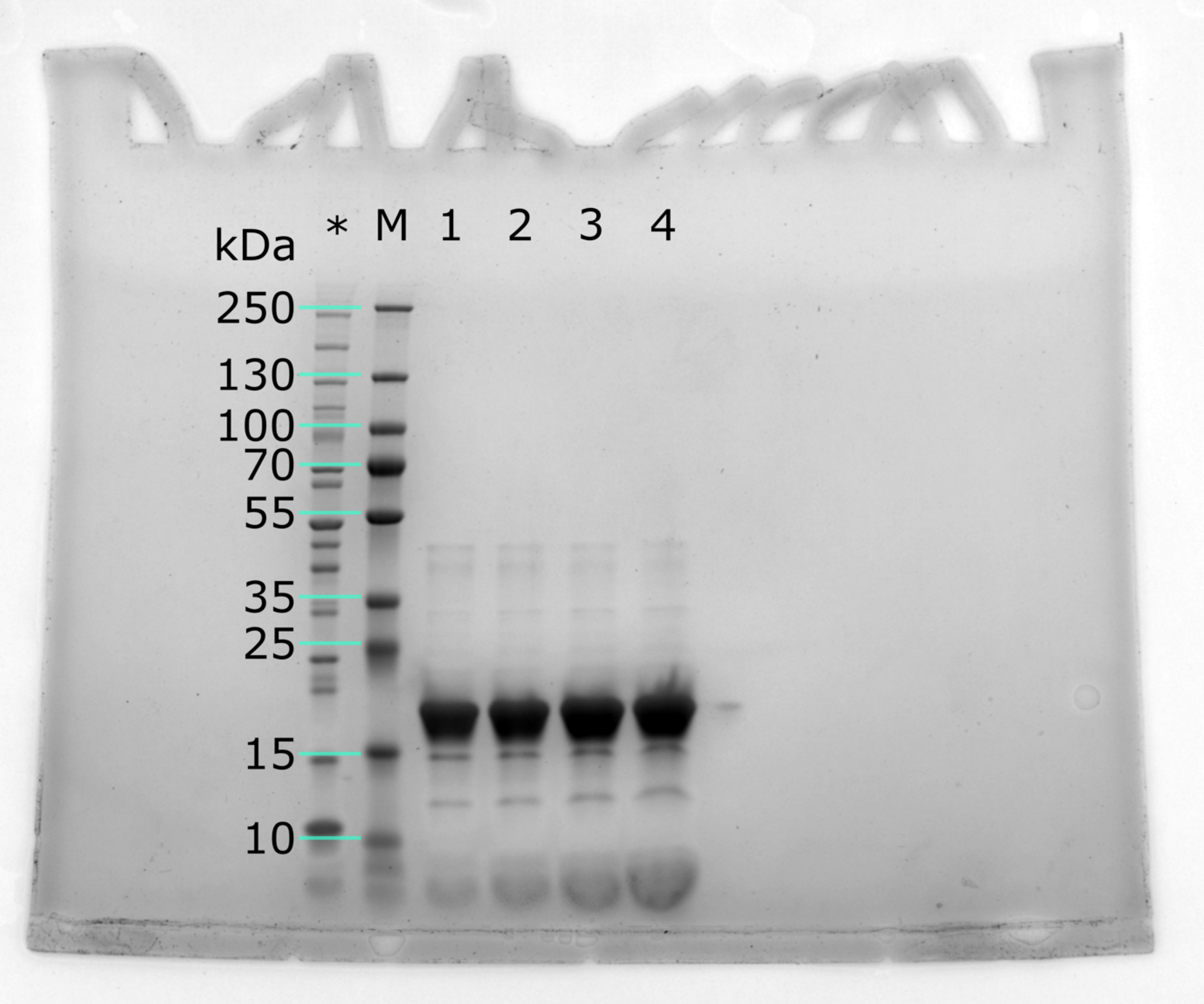


Figure S4: Uncropped SDS-PAGE of purified FGF1: 1,2,3,4 represent 12.5 ng, 25 ng, 50 ng and 100 ng FGF1 loaded in the lane. A PAGEruler unstained protein standard was also included in this gel (*), but not used for Mw estimation.

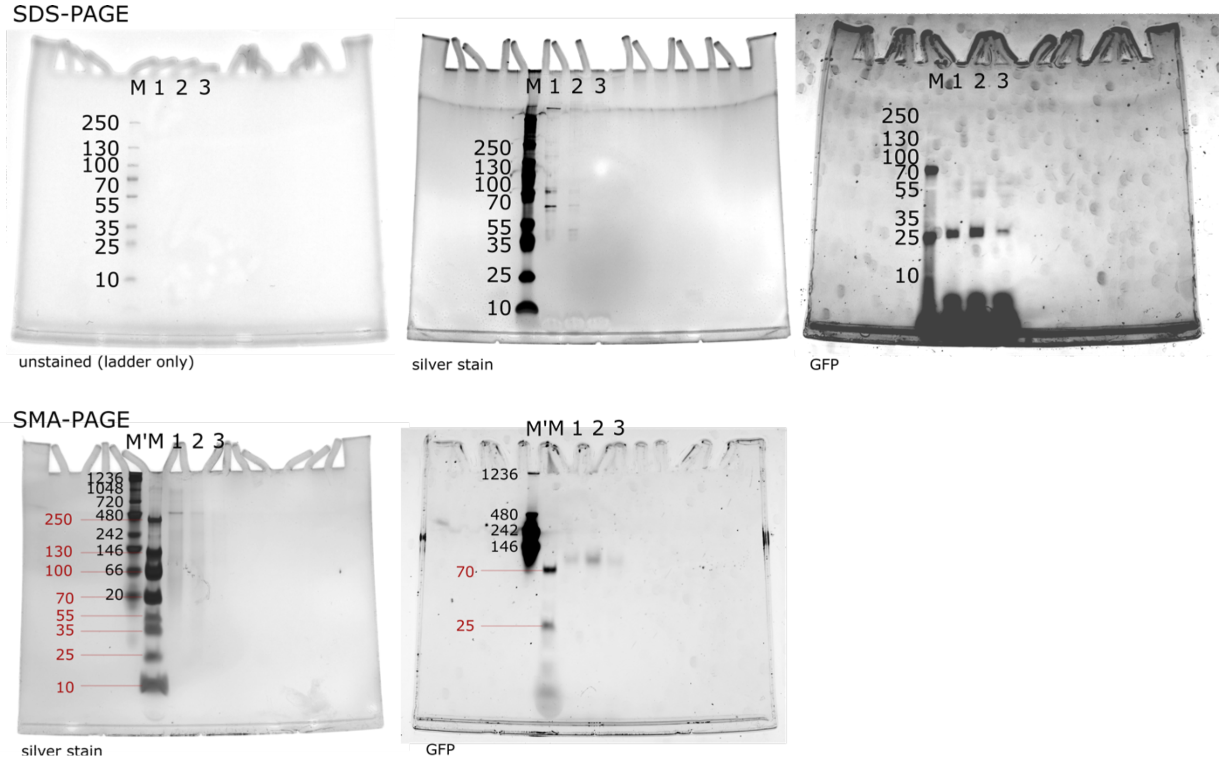


Figure S5: Uncropped PAGEs of RT112 purification, presented in Figure 1. Top row are the same SDS-PAGE imaged without stain; by silver staining; and in-gel GFP fluorescence (left to right). Bottom row are SMA-PAGEs imaged by silver stain (left) and in-gel GFP fluorescence (right). Samples 1, 2 and 3 refer to SEC peaks as shown in Figure 1. M refers to PageRuler prestained (thermo) ladder, and M' refers to NativeMark unstained (Thermo). All ladder bands visible on each imaging technique have been labelled, and ones where no clear band was visible, such as on fluorescent images, have been omitted.

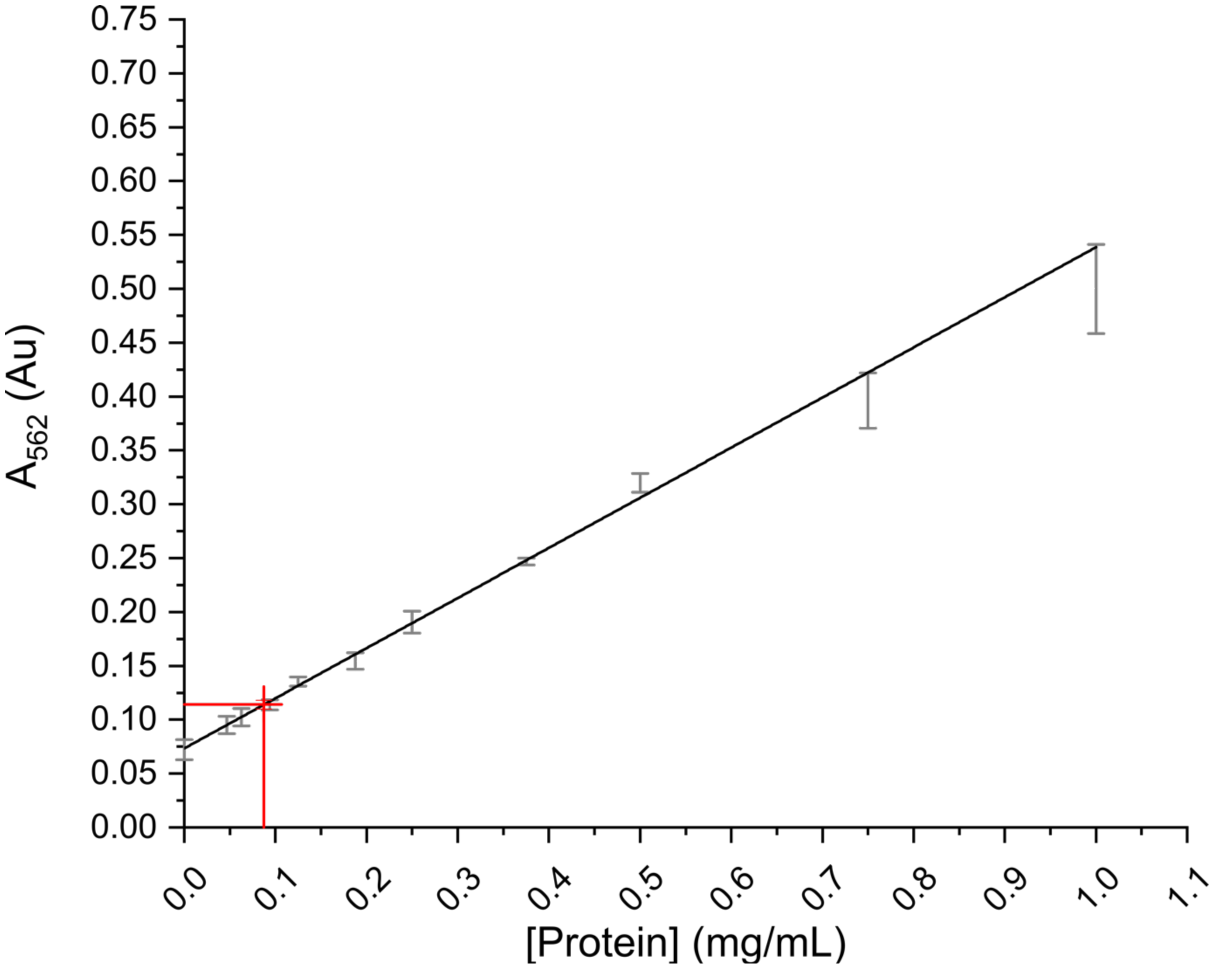


Figure S6: BCA assay of purified RT112. All standards were measured with 4 repeats (black). Test data were performed with 10 technical repeats (red). Standard line was fitted with a $t^2$ of 0.99. The specific yield was estimated based on a 1500 µL total yield from the purification producing this dataset, produced using 500 µL of ALiCE lysate.

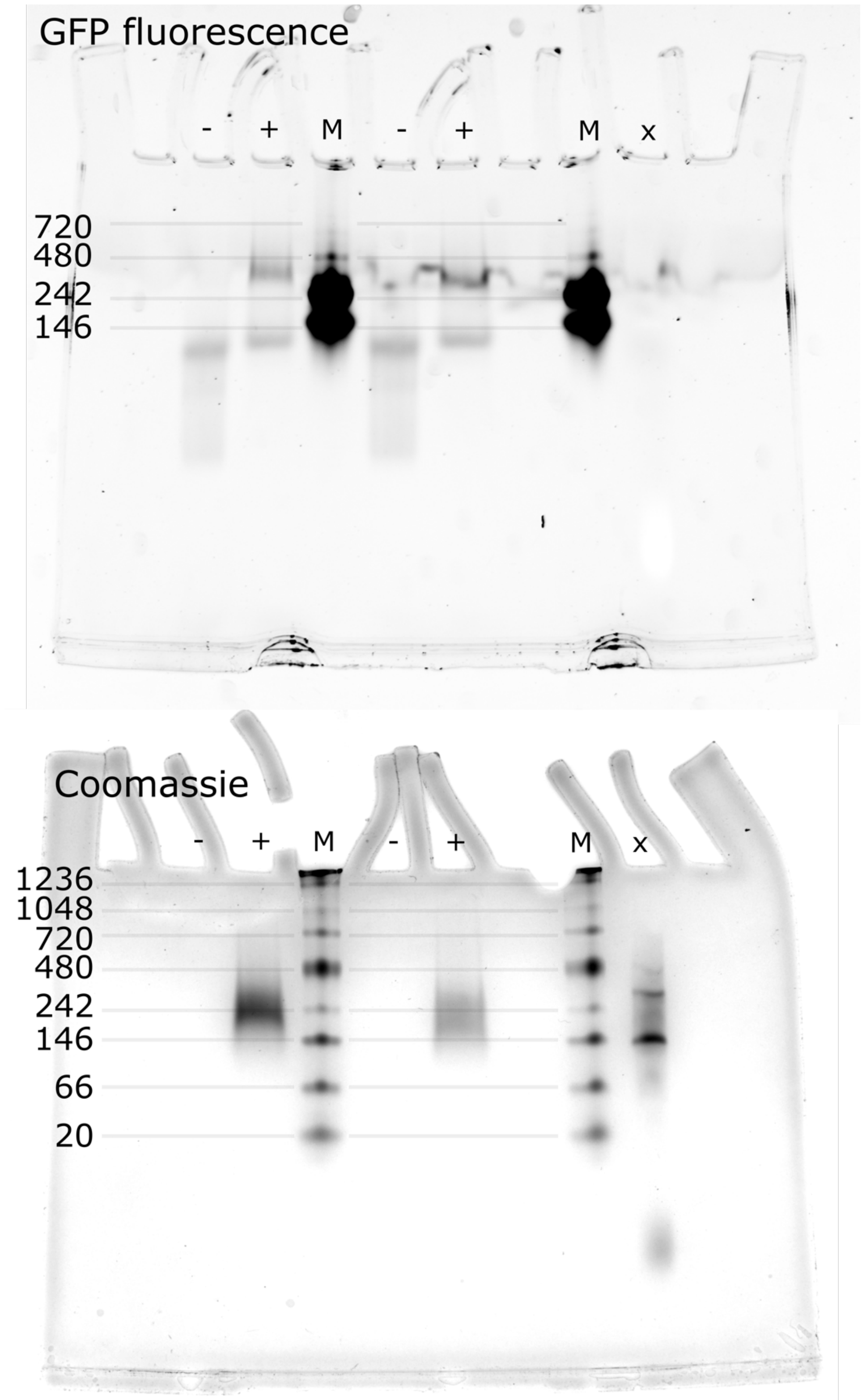


Figure S7: Uncropped SMA-PAGE of FGF1 binding assay. Top represents GFP fluorescence, bottom Coomassie staining of the same gel. Gel lanes were repeated twice to confirm result. Band visible on Coomassie stained gel in + lanes refers to unbound FGF1 (Figure S8). Lanes -,+ refer to RT112 without/with FGF1 as in Figure 3. X refers to a PageRuler unstained ladder that was stained too poorly for use in this figure.

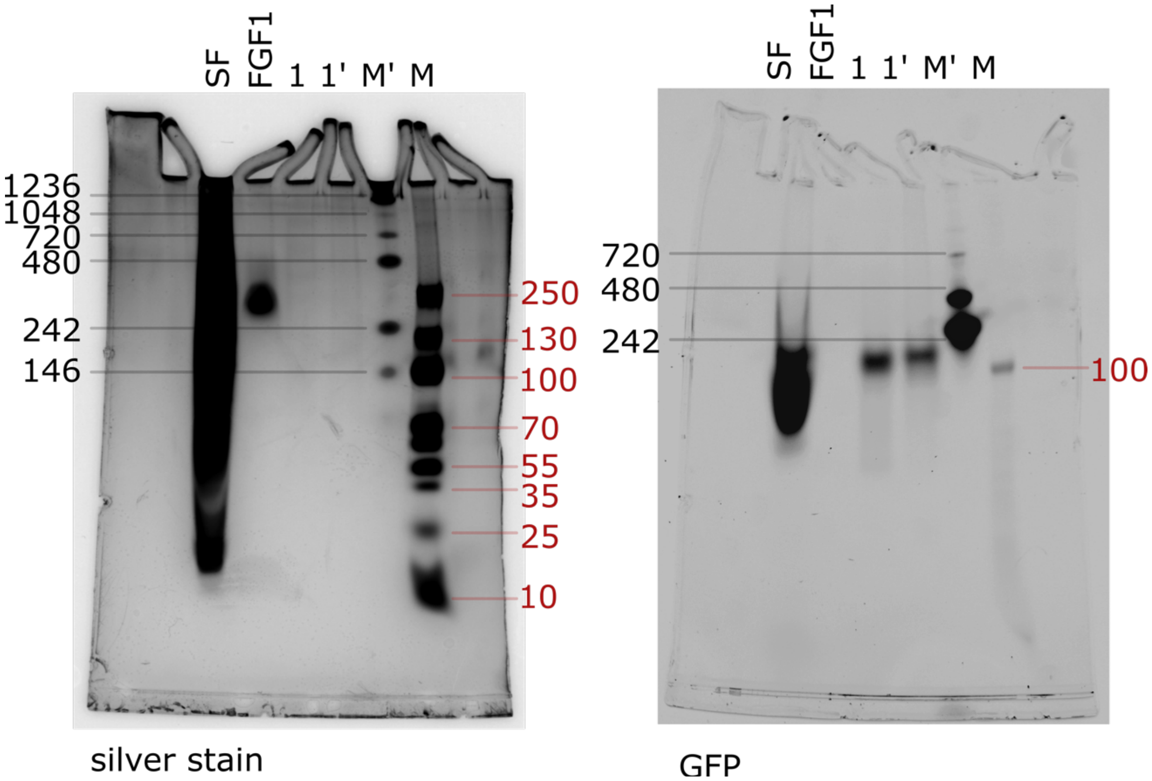


Figure S8: SMA-PAGE assessment of RT112 proteolysis. SF refers to non-membraneous proteins collected pre-solubilisation. FGF1 refers to a sample of SMA-free FGF1. 1 refers to purified RT112 from the dimeric SEC peak; 1' refers to the same with a 30 minute incubation with FGF1 prior to running. M' refers to nativemark protein ladder; M refers to PageRuler prestained ladder. Gel was imaged using in-gel GFP fluorescence and silver stain.

| Protein | Amino acid sequence |
|---|---|
| FGFR3.TACC3 RT112 variant (sequence-derived 136.4 kDa) | MPRGAPACALALCVAVAIVAGASSESLGTEQRVVGRAAEV<br>PGPEPGQQEQLVFGSGDAVELSCPPPGGGPMGPTVWVK<br>DGTGLVPSERVLVGPQRLQVLNASHEDSGAYSCRQRLTQ<br>RVLCHFSVRVTDAPSSGDDEDGEDEAEDTGVDTGAPYWT<br>RPERMDKKLLAVPAANTVRFRCPAAGNPTPSISWLKNGRE<br>FRGEHRIGGIKLRHQQWSLVMESVVPSDRGNYTCVVENK<br>FGSIRQTYTLDVLERSPHRPILQAGLPANQTAVLGSDVEFH<br>CKVYSDAQPHIQWLKHVEVNGSKVGPDGTPYVTVLKSWI<br>SESVEADVRLRLANVSERDGGEYLCRATNFIGVAEKAFWL<br>SVHGPRAAEEELVEADEAGSVCAGILSYGVGFFLFILVVAA<br>VTLCRLRSPPKKGLGSPTVHKISRFPLKRQVSLESNASMSS<br>NTPLVRIARLSSGEGPTLANVSELELPADPKWELSRARLTL<br>GKPLGEGCFGQVVMAEAIGIDKDRAAKPVTVAVKMLKDD<br>ATDKDLSDLVSEMEMMKMIGKHKNIINLLGACTQGGPLYV<br>LVEYAAKGNLREFLRARRPPGLDFSFDTCKPPEEQLTFKDL<br>VSCAYQVARGMEYLASQKCIHRDLAARNVLVTEDNVMKIA<br>DFGLARDVHNLDYFKETTNGRLPVKWMAPEALFDRVYTH<br>QSDVWSFGVLLWEIFTLGGSPYPGIPVEELFKLLKEGHRM<br>DKPANCTHDLFMIMRECWHAAPSQRPTFKQLVEDLDRVL<br>TVTSTDVKATQEENRELRSRCEELHGKNLELGKIMDRFEE<br>VVYQAMEEVQKQKELSKAEIQKVLKEKDQLTTDLNSMEKS<br>FSDLFKRFEKQKEVIEGYRKNEESLKKCVEDYLARITQEGQ<br>RYQALKAHAEEKLQLANEEIAQVRSKAQAEALALQASLRK<br>EQMRIQSLEKTVEQKTKENEELTRICDDLISKMEKIPAGL**EV**<br>**LFQGP**H**MSKGEELFTGVVPILVELDGDVNGHKFSVSGE**<br>**GEGDATYGKLTLKFICTTGKLPVPWPTLVTTFSYGVQCFS**<br>**RYPDHMKRHDFFKSAMPEGYVQERTISFKDDGNYKTR**<br>**AEVKFEGDTLVNRIELKGIDFKEDGNILGHKLEYNYNSH**<br>**NVYITADKQKNGIKANFKIRHNIEDGSVQLADHYQQNT**<br>**PIGDGPVLLPDNHYLSTQSKLSKDPNEKRDHMVLLEFV**<br>**TAAGITHGMDELYK**TAGSAAGSASIMRGSHHHHHHHH |
| Fibroblast Growth Factor 1 (FGF1)<br>Sequence-derived 18.52 kDa | MAEGEITTFTALTEKFNLPPGNYKKPKLLYCSNGGHFLRILP<br>DGTVDGTRDRSDQHIQLQLSAESVGEVYIKSTETGQYLAM<br>DTDGLLYGSQTPNEECLFLERLEENHYNTYISKKHAEKNW<br>FVGLKKNGSCKRGPRTHYGQKAILFLPLPVSSDLEHHHH<br>HH |

Table S1: Amino acid sequences of constructs used in this study. For RT112, a thrombin-cleavage site (red) links the protein to GFP (green) and the affinity tag. C228R, S249C and Y375C were introduced into RT112 to increase dimer stability.
